# Reference-free deconvolution of DNA methylation data and mediation by cell composition effects

**DOI:** 10.1101/037671

**Authors:** E. Andres Houseman, Molly L. Kile, David C. Christiani, Tan A. Ince, Karl T. Kelsey, Carmen J. Marsit

## Abstract

We propose a simple method for reference-free deconvolution that provides both proportions of putative cell types defined by their underlying methylomes, the number of these constituent cell types, as well as a method for evaluating the extent to which the underlying methylomes reflect specific types of cells. We have demonstrated these methods in an analysis of 23 Infinium data sets from 13 distinct data collection efforts; these empirical evaluations show that our algorithm can reasonably estimate the number of constituent types, return cell proportion estimates that demonstrate anticipated associations with underlying phenotypic data; and methylomes that reflect the underlying biology of constituent cell types. Thus the methodology permits an explicit quantitation of the mediation of phenotypic associations with DNA methylation by cell composition effects. Although more work is needed to investigate functional information related to estimated methylomes, our proposed method provides a novel and useful foundation for conducting DNA methylation studies on heterogeneous tissues lacking reference data.

## Introduction

In the last decade, there has been an increasing interest in epigenome-wide association studies (EWAS), which aim to investigate associations between DNA methylation and health or exposure phenotypes across the genome. Numerous publications have reported associations between DNA methylation profiled in a single tissue and disease states or exposure phenotypes. Most of these studies have used whole blood^1^ or cord blood^2-4^, but some have used other media such as buccal swabs^5^, adipose tissue^6, 7^, and placenta^8-11^.

However, most tissues are complex mosaic of cells derived from at least two and sometimes three different germ layers; endoderm, mesoderm and ectoderm that give rise to both epithelial and stromal compartments. Just the epithelial component of an organ can be composed of many cell types; for example we found that breast epithelium is composed of at least 10-12 cell types^12^ with potentially distinct DNA methylation profiles^13^. Added to this complexity are the cells in the stromal component with distinct functions, including vascular and lymphoid endothelial cells and pericytes, immune cells such as macrophages, leukocytes and lymphocytes, stromal fibroblasts, myofibroblasts, myoepithelial cells, as well as adipose cells, endocrine cells, nerve cells and other cellular and tissue elements that have different but systematically varying developmental origins. The complexity of the epigenome in normal tissues has been described in a recent analysis of 111 reference human epigenomes of human tissues^14^. Thus, because normal tissue development, individual cellular differentiation and cellular lineage determination are regulated by epigenetic mechanisms, which include chromatin alterations as well as DNA methylation^15-18^, many phenotypic associations with DNA methylation may be explained in whole or in part by systematic associations with the distribution of underlying cell types. This has been demonstrated statistically in numerous papers^19-22^ and in one notable recently published manuscript which identified and confirmed the specific cell subtype responsible for the highly replicated relationship between tobacco smoking exposure and DNA methylation of the GPR15 locus^23^. This phenomenon has led to an interest in methods for adjusting EWAS studies for cell-type heterogeneity. In *referenced-based* deconvolution methods, the distribution of cell types is obtained by projecting whole-tissue DNA methylation data onto linear spaces spanned by cell-type-specific methylation profiles for a specific set of CpGs that distinguish the cell types, so-called *differentially methylated positions* (DMPs)^19^; these methods require the existence of a reference set consisting of the cell-type specific methylation profiles, such as those that exist for blood^19, 24, 25^. However, no such reference sets exist for solid tissues of interest, such as adipose and placenta, or even tumors, thus motivating *reference-free* methods^13, 26, 27^ that seek to adjust DNA methylation associations for cell-type distribution.

Numerous cell-type deconvolution methods are currently available, many of them based on mRNA or protein expression^28^; all of them are essentially either reference-based, i.e. supervised by the pre-selection of loci known to differentiate cell types, or else reference-free, i.e. essentially unsupervised. While reference-based deconvolution methods allow for direct inference of the relationship between phenotypic variation and altered cell composition of characterized cell subtypes, reference-free approaches can provide only limited, if any, information on the types of cells contributing to the phenotypic association. In this article we propose a simple method for reference-free deconvolution that addresses this challenge and that provides both interpretable outputs – proportions of putative cell types defined by their underlying DNA methylation profiles – as well as a means for evaluating the extent to which the underlying profiles reflect specific types of cells.

Our fundamental approach is as follows: we assume an *m × n* matrix **Y** representing DNA methylation data collected for *n* subjects or specimens, each measured on an array of *m*CpG loci, and that the measured values are constrained to the unit interval [0,1], each roughly representing the fraction of methylated cytosine molecules in the given sample at a specific genomic position. This conforms to the typical *average beta* output of popular platforms such as the Infinium arrays by Illumina, Inc. (San Diego, CA), i.e. the older HumanMethylation27 (27K) platform, which interrogates 27,578 CpG loci, and the newer HumanMethylation450 (450K) platform, which interrogates 485,412 CpG loci; however, it also conforms to the results of sequencing-based platforms such as whole genome bisulfite sequencing (WGBS). In reference-based methods, the following relation is assumed to hold: **Y** = **MΩ**^T^, where **M** is a *known m × K* matrix representing *m* CpG-specific methylation states for *K* cell types and **Ω** is an *n × K* matrix representing subject-specific cell-type distributions (each row representing the cell-type proportions for a given subject, i.e. the entries of **Ω** lie within [0,1] and the rows of **Ω** sum to values less than one). Reference-free methods attempt to circumvent lack of knowledge about **M** either by using a two-stage regression analysis (e.g. the Houseman approach^27^) or else fitting a high-dimensional mixed-effects model and equating the resulting random coefficients with cell-mixture effects (i.e. the Zou approach^26^); both methods rely on a predetermined model positing associations between DNA methylation **Y** and phenotypes **X**. For example, the Houseman method posits the model **Y** = **AX**^T^ + **R**, where **X** is an *n × d* design matrix of phenotype variables and potential confounders; the *m × d* regression coefficient matrix **A** and the *m × n* error matrix **R** are both assumed to have further linear structure involving **M**, and the common variation between **A** and **R** is assumed to represent systematic association with cell type distribution. However, results of this approach are somewhat influenced by the choice of the dimension of the linear subspace of [**A, R**] representing the common variance induced by **M**^20^; consequently there has been recent concern that the method may over-adjust for cell distribution. A similar problem exists with the Zou approach, which models the phenotype as a linear function of DNA methylation, and in which the choice of a tuning parameter can influence the extent to which phenotypic associations are putatively explained by heterogeneity in underlying cell types. Here, we propose that a variant of non-negative matrix factorization be used to decompose **Y** as **Y** = **MΩ**^T^, where the entries of the unknown matrices **M** and **Ω** are constrained to lie in the unit interval and the rows of **Ω** are constrained to sum to a value less than or equal to one. This approach is similar to existing approaches for estimating the proportion of normal tissue cells in a tumor sample or otherwise deconvolving mixtures of cells^29-33^. Additionally, this factorization conforms to the biological assumption that DNA methylation measurements **Y**, regardless of associated metadata **X**, ultimately arise as linear combinations of constituent methylomes, as we have previously argued^20^. However, such constrained factorizations can be computationally intensive, and it is still necessary to specify the number *K* of assumed cell types, so in Supplementary Section S1 we propose a fast approximation that facilitates resampling, which is the basis of our method for determining *K*, described in Section S2. Note that *K* = 1 corresponds to the case where there are no relevant constituent cell types, which should be true for relatively pure media. If associations remain between **Ω** and **X**, i.e. if the associations between **X** and **Y** factor through the decomposition **Y** = **MΩ**^T^, then these associations are potentially explained by systematic changes in cell composition. Evidence for mediation of associations by cell type is substantially strengthened if the methylomes represented by **M** map to biological processes that correspond to distinct populations of cells. To that end, we propose a simple companion analytical procedure for the interpretation of the methylomes represented by **M**. Denote each row of **M** (corresponding to one CpG) as the *K × 1* vector **μ***_j_*, *j*∈ {1,…,*m*}. CpG loci that most differentiate the *K* putative cell types will tend to have distinct values within **μ***_j_*; thus high values of the row-variance 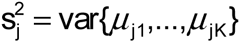 should correspond to CpGs that are most relevant to the biological distinctions among the *K* cell types, and this can be tested with auxiliary annotation data. Figure 1 illustrates our approach.

**Figure 1.**
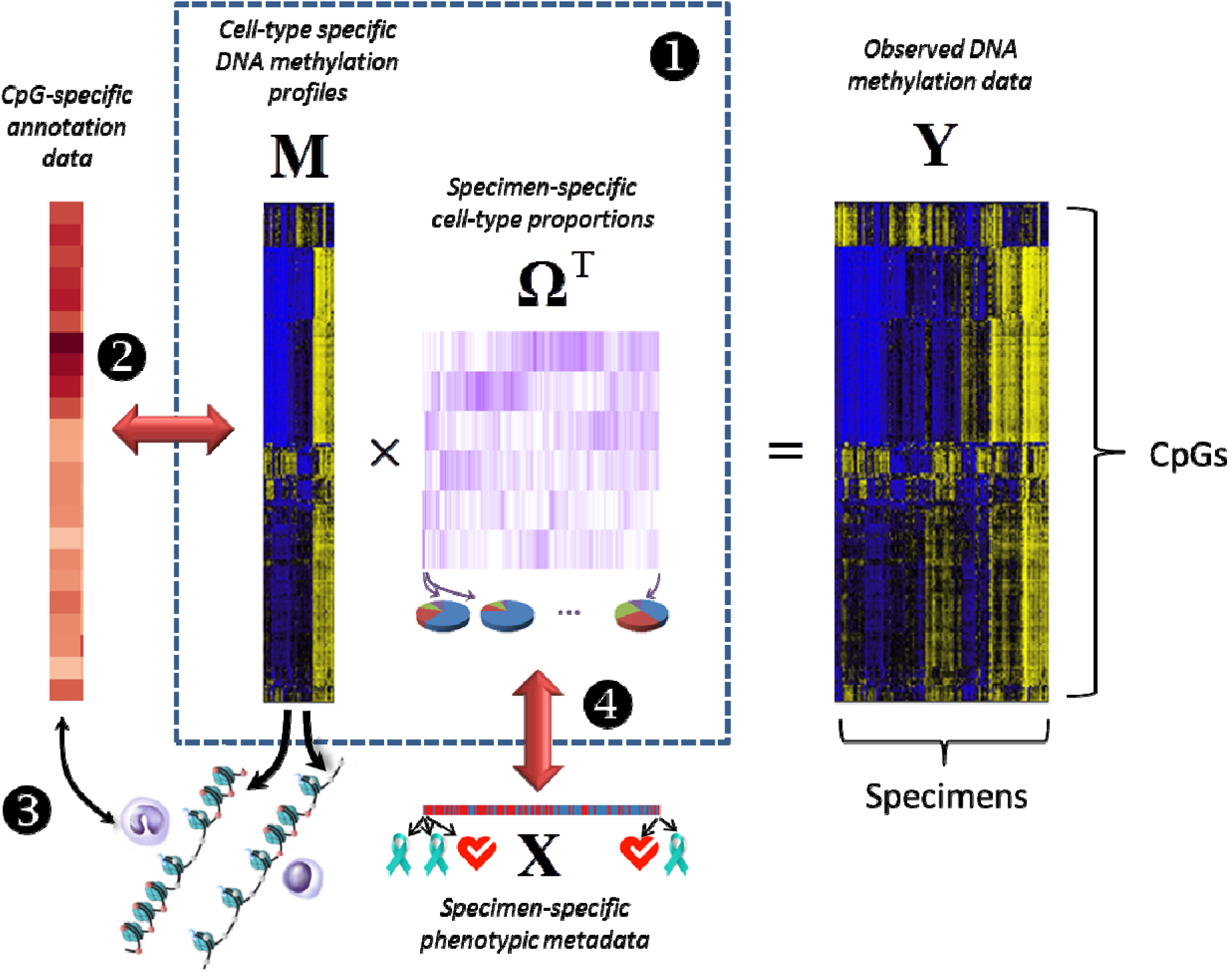
If associations between DNA methylation data **Y** and phenotypic metadata **X** factor through the decomposition **Y** = **MΩ**^T^, and the data in **M** serve to distinguish cell types by their associations with relevant annotation data, then associations between **X** and **Y** are explained in whole or in part by differences in the distribution of constituent cell types.

We demonstrate these methods in analyses of 23 genome-scale DNA methylation data sets from 13 distinct data collection efforts, including four blood data sets, several breast tumor data sets (including data from The Cancer Genome Atlas, *TCGA*), vascular and liver tissues, sperm, and four separate media collected on the same population, Bangladeshi neonates, including placenta. In addition, we leverage data derived from The Roadmap Epigenomics Project, demonstrating their utility in addressing the biological relevance of fitted methylomes **M**.

## Results

To test our proposed approach, we analyzed 23 DNA methylation datasets from 13 distinct studies, each set of DNA methylation measurements obtained via the Infinium 27K or 450K platform. Four blood data sets^3, 22, 34, 35^ were included as positive controls (given the existing reference data and known heterogeneity), each collected in the context of an epidemiologic study, and each assumed to exhibit heterogeneity in cell type as previously described^3, 19, 22^. Sperm^36^ and isolated vascular tissues^37^ were included as negative controls, presumed to represent relative homogeneity in terms of constituent cell types. Note that four datasets arose from one study on arsenic exposure in Bangladeshi neonates^3, 9^, in which four separate tissues were obtained from the same individuals. Also included were arterial tissue^38^, liver tissue^39^, and data from cancer data sets^40-43^, including breast tissues from *TCGA^44^.* Table 1 lists the data sets, their sources and their short descriptions. Figure S3.1 shows the results of hierarchical clustering applied to 26,476 CpG sites common across the datasets (Manhattan distances based on mean methylation for the data set, clustering based on Ward’s method implemented as *Ward.D* in R version 3.2.2). Figure S3.2 summarizes the number of CpGs analyzed for each data set, by fraction of samples observed for each CpG. Note the strong clustering of data sets by type of media. The ordering of data sets in Table 1 and many subsequent figures are based on the clusters shown in Figure S3.1.

**Table 1.**
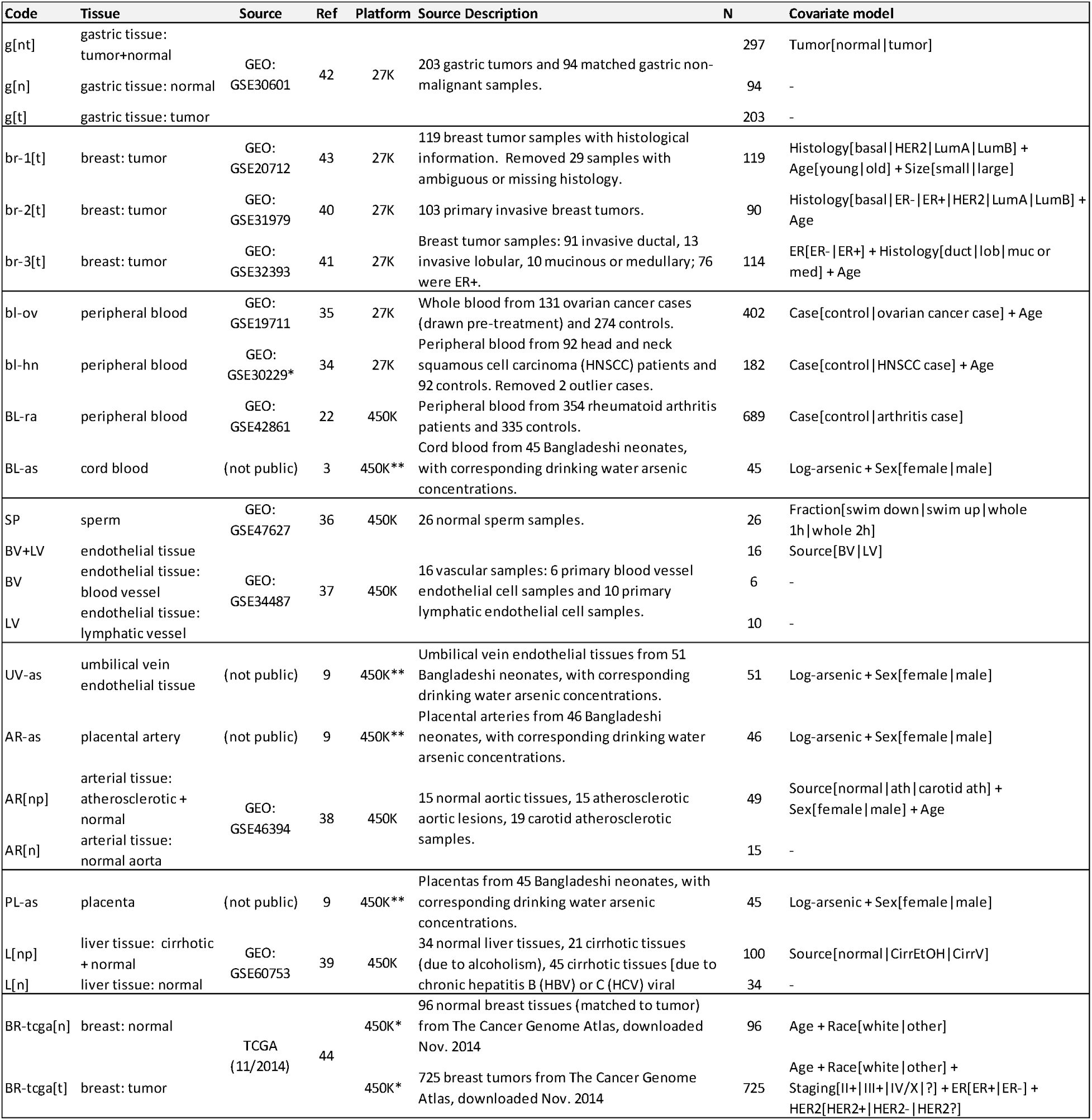
Summary of Datasets

### Estimated Numbers of Cell Types

Using the method described in Supplementary Section S2 with 25 iterations, for each data set we found the decomposition **Y** = **MΩ**^T^, for values of *K* varying from 2 to either *K*_max_ = 10 or the maximum possible given the sample size (*K*_max_ = 8 for *BV+LV*, 2 for *BV* and *LV*, 7 for *AR[n]*). We then used our bootstrap approach (Supplementary Section S3) for determining the number 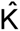 of classes for each data set, displayed in Figure 2a, which demonstrates heterogeneity in the number of classes 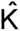 estimated. 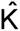 ≥ 3 for blood data sets (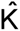 = 3 for the cord blood data set *BL-as*, larger for the other three peripheral blood datasets). For breast tissues (both tumor and normal) 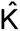 was typically large. 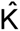 ≥ 3 for the artery and liver data sets having three distinct sources each (*AR[np]* and *L[np]*). 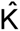 = 1 for the pure blood and lymphatic vessel data sets (*BV* and *LV*); 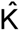 = 2 for other vessel data sets consisting of normal tissue (*AR[n]*, *AR-as, BV+LV, UV-as)*, for the normal liver data set *L[n]*, for sperm (*SP*) and for placenta (*PL-as*). We remark that 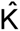 was typically lower for datasets that were more likely to be comprised of homogeneous tissues. We also remark that our proposed method of selecting 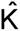 is based on minimizing a bootstrapped deviance statistic, and that the variation of this statistic across values of *K* can be informative. For example, with the *BL-ra* dataset, the deviance dropped precipitously from *K* = 1 to *K* = 3, while for the sperm data set the deviance remained flat from *K* = 1 to *K* = 6 before rapidly increasing (Figure 2b).

**Figure 2.**
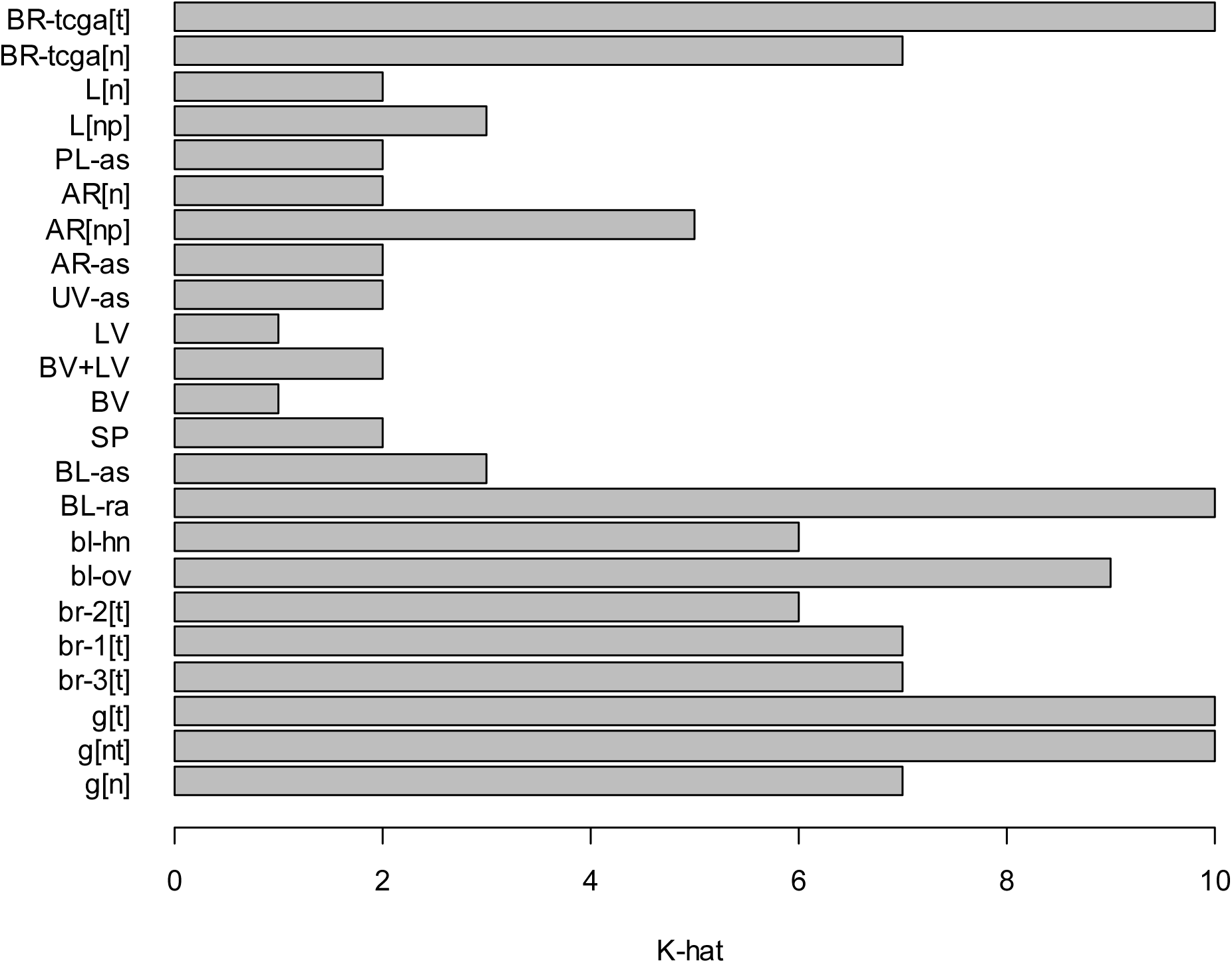
(A) Estimated number 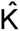 of classes for each data set. (B) Bootstrapped deviance profiles for four selected data sets, along with mean deviance, median deviance, and quartiles for each value of *K*.

### Associations with Phenotypic Metadata

To examine the associations between **Ω** and various metadata associated with the subjects/specimens in the corresponding study, we fit a quasi-binomial model for each row of **Ω**; Table 1 provides the covariate model **X** used for each data set. As described below and in detail in Supplementary Section S4, to circumvent dependence of results on the choice of *K*, we examined associations over the range *K* ∈ {1,…, *K*_max_}, using a permutation test (1000 permutations) for inference on each covariate. Table 2 provides a summary of permutation test results. As shown in Table 2, cell mixture proportions **Ω** were typically significantly associated with major phenotypes of interest and occasionally with age (e.g. *bl-hn* and *BR-tcga[t]*); the exception was the sperm dataset, for which **Ω** was not significantly associated with fraction. Note in particular that for breast tumors, ER status variables (or histology variables incorporating ER status) were significantly associated with **Ω**. Sex was typically not significantly associated with **Ω**. As shown in Figure 2, the associations between **Ω** and phenotype could be quite striking (e.g. rheumatoid arthritis status) or completely lacking (e.g. sperm fraction). Figure 3a shows clustering heatmap of **Ω** (*K* = 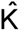 = 10) for *BL-ra*, one of the positive control blood data sets, with annotation track showing the associated phenotype rheumatoid arthritis case/control status. Figure 3b shows a similar clustering heatmap for the negative control *SP* (sperm, *K* = 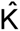 = 2), along with the associated phenotype, specimen fraction. Other clustering heatmaps are provided supplementary files.

**Table 2.** Inference with Phenotypic Metadata

| Data Set | Permutaton P-values |
| --- | --- |
| g[nt] | Tumor<0.001 |
| br-1[t] | Histology<0.001; Age=0.059; Size=0.016 |
| br-2[t] | Histology<0.001; Age=0.06 |
| br-3[t] | ER<0.001; Histology=0.295; Age=0.008; BSC=0.297 |
| bl-ov | Case<0.001; Age=0.999 |
| bl-hn | Case<0.001; Age<0.001 |
| BL-ra | Case<0.001 |
| BL-as | Log-arsenic<0.001; Sex=0.263 |
| SP | Fraction=0.994 |
| BV+LV | Source=0.013 |
| UV-as | Log-arsenic=0.515; Sex=0.962 |
| AR-as | Log-arsenic=0.285; Sex=0.505 |
| AR[np] | Source<0.001; Sex=0.043; Age=0.377 |
| PL-as | Log-arsenic=0.006; Sex=0.451 |
| L[np] | Source<0.001 |
| BR-tcga[n] | Age=0.089; Race=0.153 |
| BR-tcga[t] | Age<0.001; Race<0.001; Staging=0.013; ER<0.001; HER2<0.001 |

**Figure 3.**
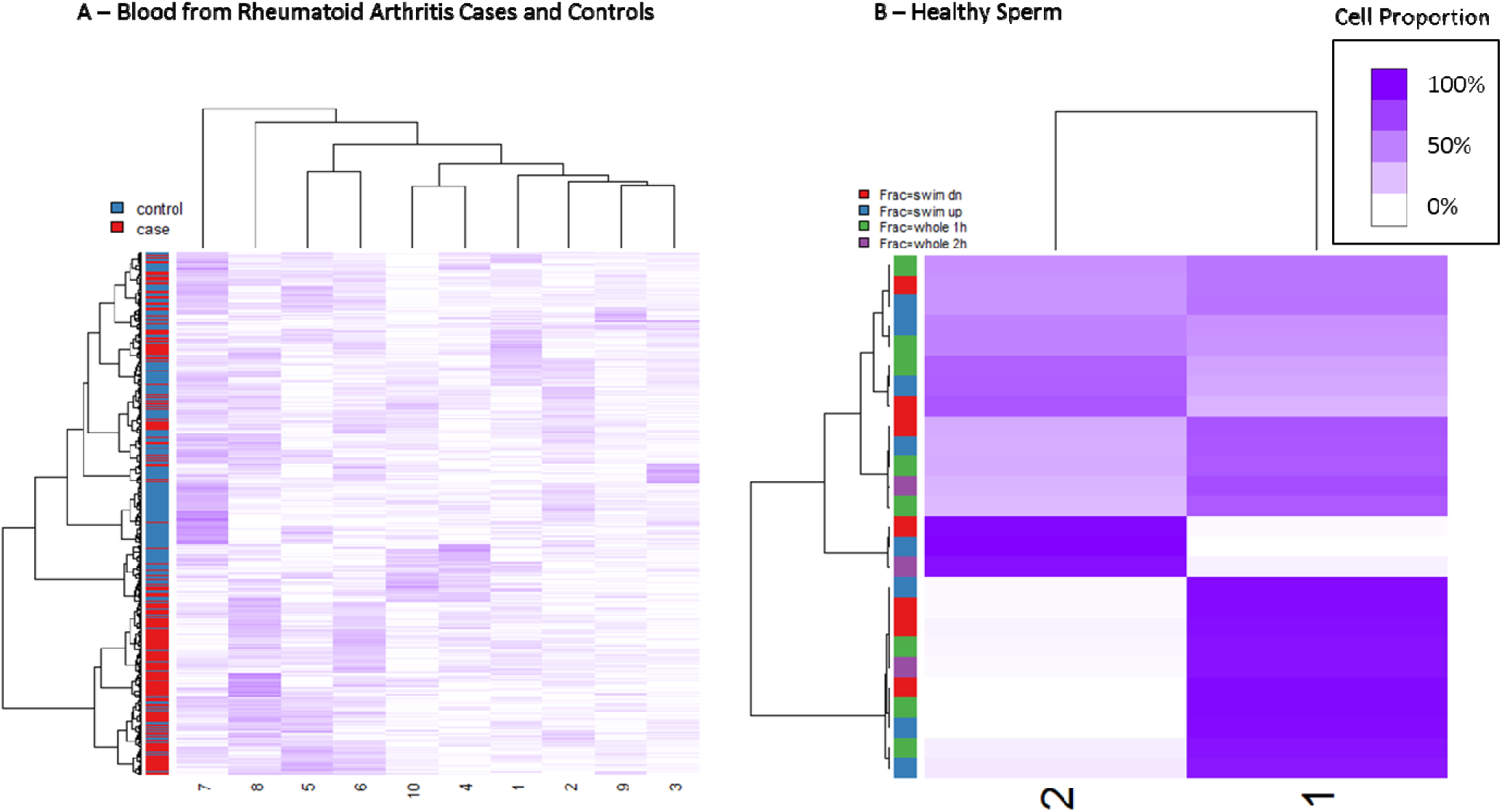
Clustering heatmaps of cell proportion matrix **Ω** for two data sets; purple intensity indicates cell proportion. (A) Blood from rheumatoid arthritis cases and controls (*BL-ra*, *K* = 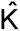 = 10). (B) Sperm (*SP*, *K* = 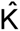 = 2).

We also considered the effect of **Ω** on CpG-specific associations of DNA methylation with **X**. As described below and in detail in Section S5, we computed regression coefficients for logit-methylation (i.e. M-values) upon [**X, Ω**] for *K* ∈ {1,…, *K*_max_} (for *K* = 1 the covariate model was simply **X**); for **Ω** and for each covariate, we then used the resulting nominal p-values to estimate the proportion *π*_0_ of null associations. For demographic variables (age, sex, race), Figure S5.1 illustrates the value of *π*_0_ for the overall association of **Ω** with DNA methylation ( *K* = *K*^*^ = max(2, 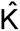)). For demographic variables, Figure S5.2 provides a comparison of *π*_0_ from the *K* = 1 model with *π*_0_ from the *K* = *K*^*^ model. Figure 4 displays a similar comparison for other variables. These figures demonstrate that adjustment by **Ω** very often resulted in higher values of *π*_0_, the estimated proportion of null associations. Exceptions were age in *bl-hn* and sex in *AR[np]* and *AR-as*, where adjustment by **Ω** reduced *π*_0_ (Figure S5.2). On the other hand, the proportion of null associations with **Ω** was typically low: except for the homogeneous tissue datasets *SP* and *BL* + *LV*, *π*_0_ was less than 0.2; of the others, except for the arsenic-exposure data sets *UV-as*, *AR-as*, and *PL-as*, *π*_0_ was extremely close to zero.

**Figure 4.**
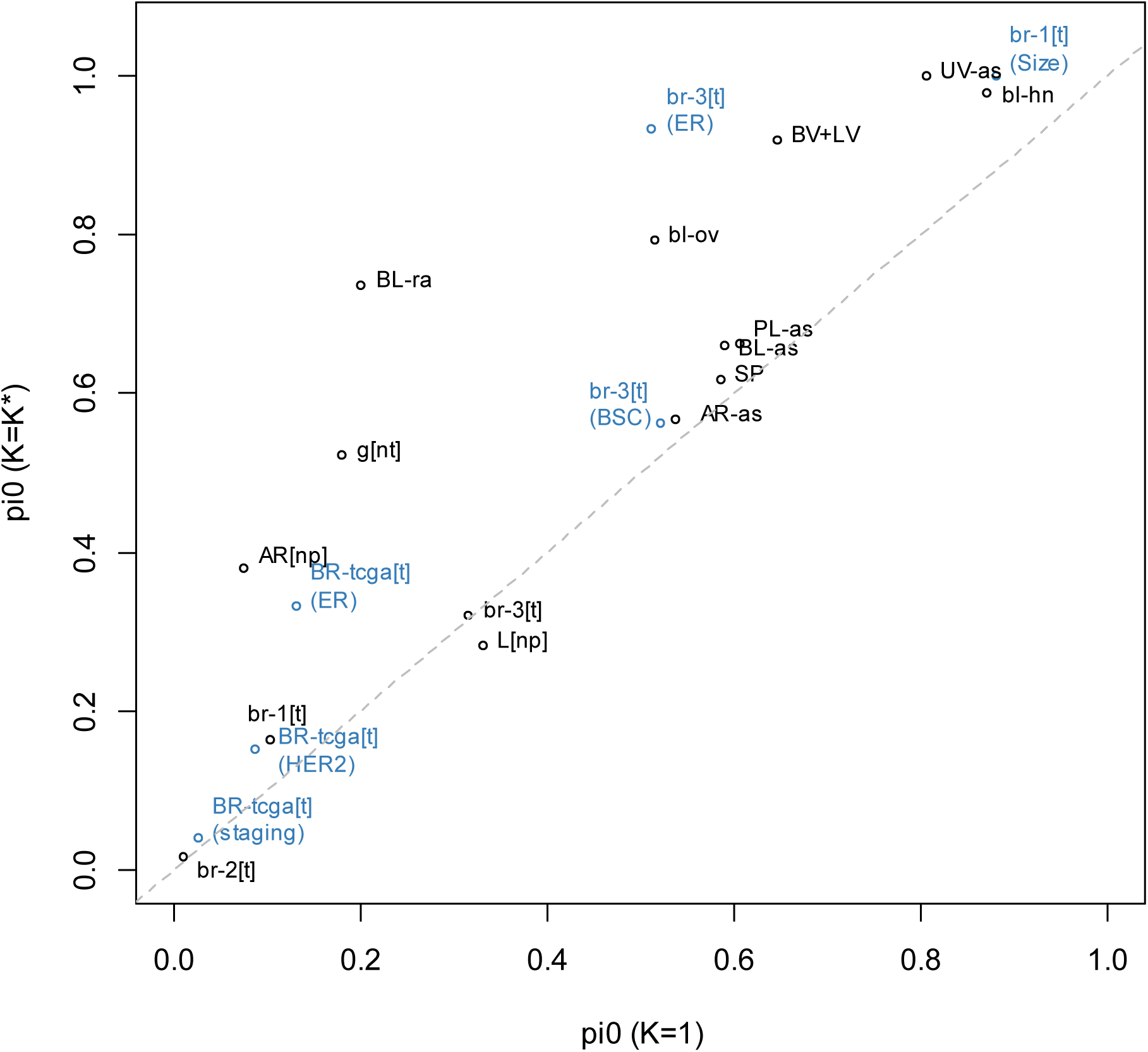
Comparison of *π*_0_ (proportion of null association CpGs) from the *K* = 1 model with *π*_0_ from the *K* = *K*^*^ model; only non-demographic variables are shown.

### Interpretation of Putative Cell Types

We examined the biological relevance of resulting matrices **M** in several different ways. First, for each data set, we computed row-variances 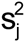 (as described above) both for *K* = 2 and for *K*^*^ = max(2, 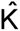). For each of these two values of *K*, we classified each CpG *j* ∈ {1,…, m} by whether its row-variance 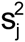 lay above the 75^th^ percentile q_0.75_(s^2^), reasoning that these CpGs could be considered as important distinguishers of cell type. Next, we obtained a list of DMPs for differentiating distinct major types of leukocytes (*Blood DMPs*), and another list of CpGs mapped to genes considered Polycomb Group proteins (*PcG loci*), the construction of both lists described in detail in Supplementary Section S6. For each data set we computed the odds ratio for the association of high row-variance 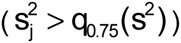 with DMP set membership (*Blood DMPs* or *PcG loci*), using Fisher’s exact test to compute the corresponding p-values. Odds ratios are depicted in Figure 5, with log_10_ p-values given in Table S6.1. *Blood DMP* status showed the highest associations in blood data sets, although also somewhat high associations in *L[np]* and *AR[np]* (data sets having tissues with potentially inflammatory components to pathology). Data sets with tumors *(BR-tcga[t], br-1[t], br-2[t], br-3[t], g[nt]*, and *g[t])* showed high association of *PcG loci* with cell-type distinguishing CpGs, but so did the data set with normal gastric tissue, *g[n]*. As shown in Figure S6.1, the *Bilenky* DMPs based on breast tissue showed the highest association with cell-type distinguishing CpGs in the data sets with breast tissue, although associations were also high in *L[np], AR[n]*, and *AR[np]*. As shown in Figure S6.2, the *REMC* DMPs, based on comparison of ectodermal/mesodermal/endodermal distinctions among embryonic stem cells, showed relatively weak (or negative) associations with cell-type distinguishing CpGs for all datasets.

**Figure 5.**
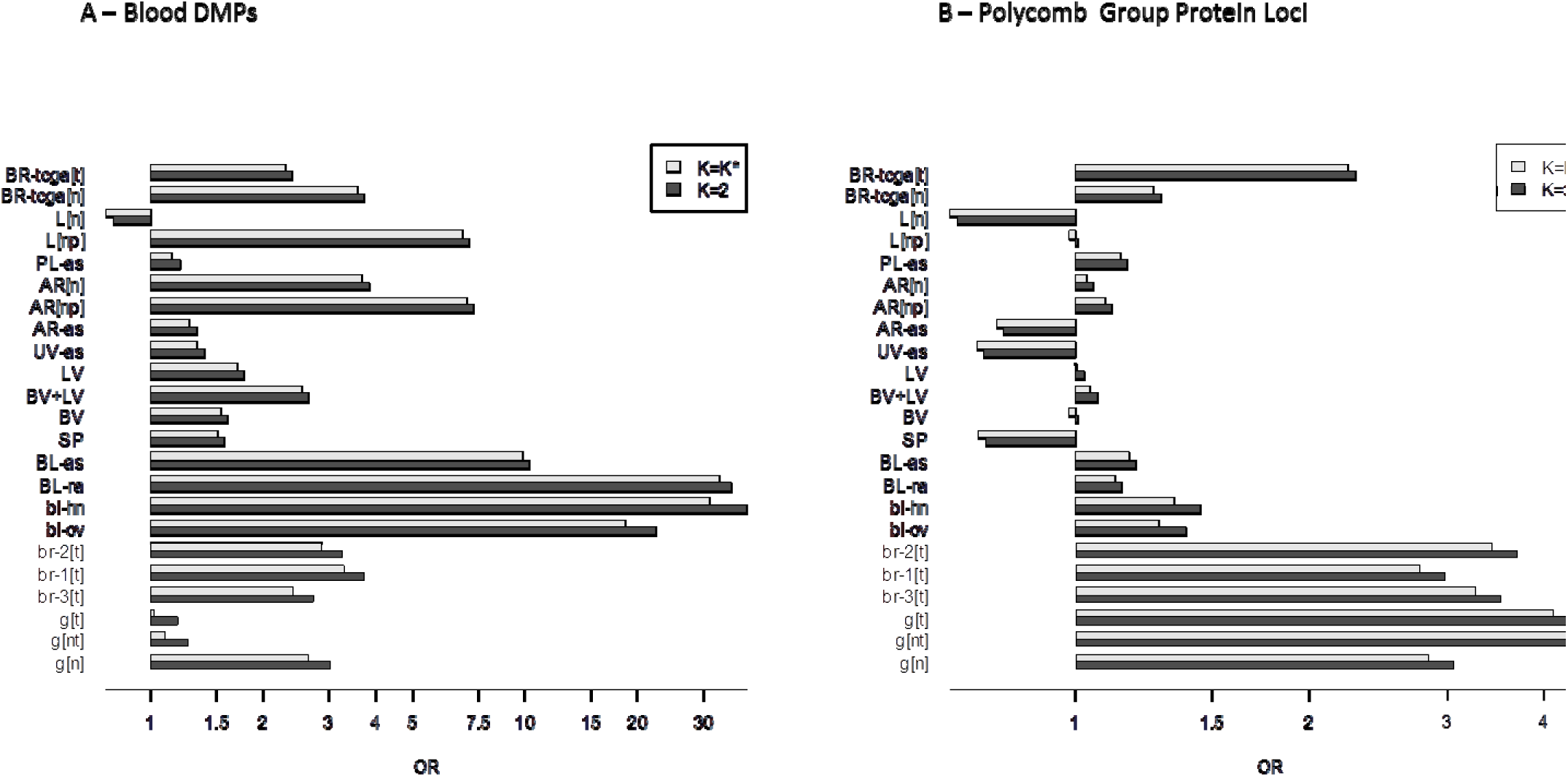
Gene-set odds ratios, showing the association of gene set membership with the set
of CpGs whose values are highly variable across fitted methylomes 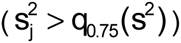. (A) Blood DMRs. (B) CpGs mapped to polycomb group protein genes.

We also developed a novel approach based on WGBS data from the Roadmap Epigenomics Project for 24 primary tissues. For each sample, we obtained the 470,909 CpGs overlapping with CpGs from either Infinium array (and having fewer than 3 missing values), clustering the tissue samples based on the 15,000 most variable of these CpGs (Manhattan distance metric with Ward’s method of clustering). The resulting dendrogram, shown in Figure S7.1, demonstrates substantial clustering along general tissue type. We also applied our deconvolution algorithm to these 24 tissue samples (*K* = 6), with results shown in Figure S7.2; note that the deconvolution of these tissues resulted in constituent cell types that roughly aligned with anticipated anatomical associations, e.g. tissues with substantial smooth or skeletal muscle mapped to one cell type, tissues with a substantial lymphoid/immune component mapped to another, and central nervous tissues map to yet another. We reasoned that *similar* tissue types would differ principally in the proportion of underlying normal constituent cell types, and thus provide information on cell-type heterogeneity underlying other tissues of similar type. Consequently, we selected the tissue pairs corresponding to the 25 smallest Manhattan distances (as calculated for the clustering in Figure S7.1), with pairs illustrated as network edges in Figure S7.3. Due to small numbers of DMPs (10 or fewer) we excluded two pairs; for each of the remaining 23 pairs, we identified, among the 15,000 CpGs most variable across all 24 tissue types, those CpGs that differed in methylation fraction by greater than 0.70 between the two samples; we considered these CpGs to be Infinium-specific DMPs for tissue-specific heterogeneity. Using these 23 sets of DMPs, we conducted a gene-set analysis as described in the previous paragraph. The clustering heatmap in Figure 6 presents the odds ratios for the 450K data with *K*^*^ = max(2, 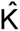); the heatmap in Figure S6.4 presents the odds ratios for the 27K data with *K*^*^ = max(2, 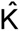), and the odds ratios for *K* = 2 are given in Figures S7.5 and S7.6. Corresponding p-values are given in Tables S7.1, S7.2 and S7.3. Note that we excluded additional pairs from the 27K array analysis due to small DMP overlap with the 27K array. As shown in Figure 6, positions that distinguished immune-related tissues (CD34+ hematopoietic stem cells vs. thymus or spleen) were highly associated with CpGs distinguishing cell types in the two 450K blood data sets, as well as in the mixed liver tissue dataset *L[np]* and the mixed arterial dataset *AR[np]*, consistent with the findings demonstrated in Figure 5a. In the arterial data sets *AR[n]* and *AR[np]*, the normal breast data set *BR-tcga[n]* and to some extent the normal mixed vessel data set *BV+LV*, high associations were found for CpGs that distinguished smooth muscle content (aorta vs. psoas muscle, heart atrium vs. ventricle, heart atrium vs. esophagus). Interestingly, *AR[n], AR[np]*, and *BR-tcga[n]* displayed associations with CpGs distinguishing lung and esophagus, potentially an epithelial cell comparison (although potentially also representing a distinction in smooth muscle content). All other positive associations were relatively weak. Strong negative associations with CpGs that distinguished right atrium from left ventricle were observed for *L[n], SP, UV-as*, and *AR-as*, although these results may be driven by small numbers of CpGs (see p-values in Table S7.2). Patterns were similar for *K* = 2 (Figure S7.5). Patterns were similar in 27K blood data sets; additionally, the normal gastric data set g[n] displayed high association with DMPs distinguishing Roadmap stomach tissues (Figure S7.4). Interestingly, *L[n]* was the only dataset displaying mostly negative (though weak) associations.

**Figure 6.**
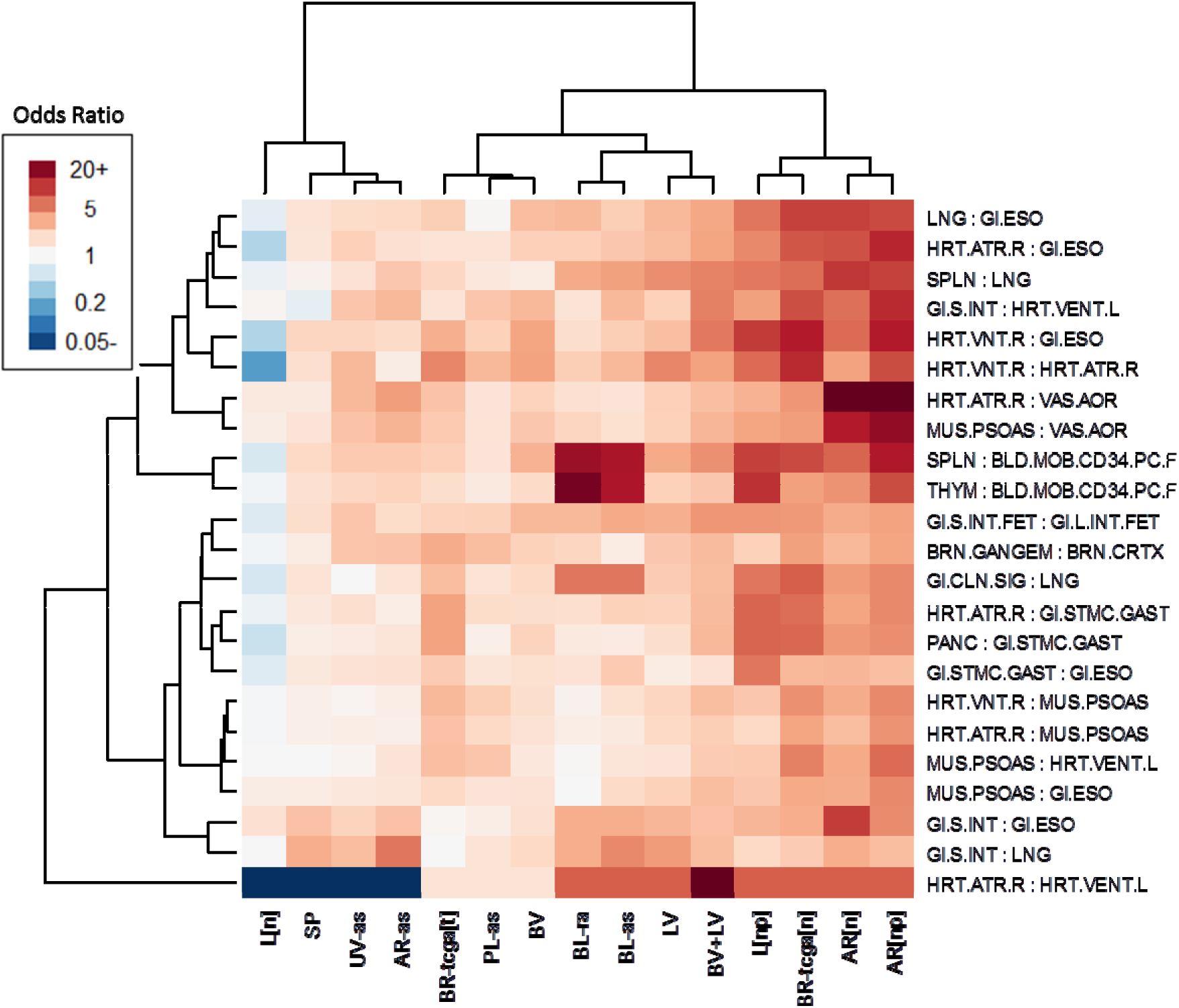
Gene-set odds ratios for 450K data sets, showing association of sets of DMPs distinguishing various Roadmap Epigenomics WGBS specimens with the set of CpGs whose values are highly variable across fitted methylomes 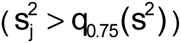.

### Additional Analyses on 450K Blood Datasets

We analyzed four blood data sets as positive controls, with the expectation that the resulting cell-type proportions **Ω** would show substantial associations with **X**. Practically, considering that reference data sets exist for blood, estimation of associations between phenotypic metadata and major types of leukocytes would typically employ the reference-based estimation of **Ω** rather than the essentially unsupervised approach we have proposed. Two additional avenues of investigation emerge: (1) the extent to which the reference-based and reference-free approaches are consistent in their results; and (2) the extent to which the unsupervised approach may provide additional information on immune response and inflammation (as represented by distributions of leukocytes including their various activation states) beyond associations with simply the major types of leukocytes, i.e. those existing in currently available reference sets. To this end, we further analyzed the two 450K blood data sets, *BL-ra* and *BL*as, estimating for each data set two sets of cell-type proportion matrices (*K* = 7): **Ω**_0_ (reference-based) and **Ω**_1_ (reference-free). We used a common set of DMPs for each estimation procedure, with details provided in Supplementary Section S8. Note that for the reference-based approach, we fit 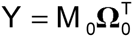 with essentially known **M**_0_, while for the reference-free approach, we estimated **M**_1_ in the context of fitting 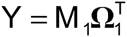. We note that, in general, we did not anticipate **Ω**_0_ and **Ω**_1_ to be equal. The reason is that the unsupervised, reference-free approach will find only the major axes of variation within a given data set, not necessarily all relevant distinctions of major cell types. For example, if a data set consists of only two distinct immune profiles (with very little variation among the subjects sharing a profile), then the reference-free approach will typically find only two cell types, those corresponding to each profile. However, **M**_0_ and **M**_1_ should be related to by a mixing matrix **Ψ** that reassigns the “correct” cell types to the unsupervised decomposition, i.e. **M**_1_ = **M**_0_**Ψ**^T^. It follows that **Ω**_0_ ≈ **Ω**_1_**Ψ**, thus phenotypic associations with **Ω**_0_ should match those with **Ω**_1_**Ψ**. Figures S8.1 and S8.2 depict the mixing matrices **Ψ**, obtained by constrained projection, for *BL-ra* and *BL-as*, respectively. Though our essentially unsupervised approach resulted in cell proportion estimates **Ω**_1_ quite distinct from **Ω**_0_, for both *BL-ra* and *BL-as* the reference-based solution **Ω**_0_ was nevertheless reasonably similar to a linear combination **Ω**_1_**Ψ** of the unsupervised solution **Ω**_1_ (Figures S8.3 and S8.4). Phenotypic associations with the “re-mixed” **Ψ**_1_**Ψ** cell proportion estimates were remarkably similar to associations with the reference-based solution **Ω**_0_ (Figures S8.5 and S8.6), with only one notable reversal: in the *BL-ra* dataset, the relative magnitudes of CD4+ and CD8+ coefficients were reversed, but all were still significantly and negatively associated with rheumatoid arthritis status.

It follows that if **M**_1_ contains information on immune function not readily apparent from **M**_0_, then the information should be evident in the residual matrix **M**_1_ – **M**_0_**Ψ**^T^. In fact, a number of CpGs still showed substantial heterogeneity among the residual methylomes **M**_1_ in comparison with the residual methylomes **M**_1_ – **M**_0_**Ψ**^T^ (Figures S8.7 and S8.8). The residual methylomes **M**_1_ – **M**_0_**Ψ**^T^, which reflect residual epigenetic information suggestive of cell-type heterogeneity but unaccounted for by the reference methylomes **M**_0_, displayed substantially diminished association with the DMPs based on Roadmap WGBS data, compared with the unadjusted methylomes **M**_1_ (Figure S8.9 vs Figure S8.10). However, with loci mapped to genes known to reflect immune activation or regulation, both **M**_1_ and **M**_1_ – **M**_0_**Ψ**^T^ typically displayed similar heterogeneity across constituent methylomes (Figures S8.11 and S8.12). In particular, they identified two strongly significant processes in the rheumatoid arthritis data set, *Th1 & Th2 differentiation* and *T-Cell Polarization* (Figure S8.13), while for arsenic exposure in Bangladeshi neonates, they identified one strongly significant process, *Regulators of T-Cell Activation* (Figure S8.14). These results were somewhat dissimilar from results obtained using the reference-based cell proportions **Ω**_0_ in *limma* to adjust for cell type (Figures S8.15 through S8.17). In particular, the limma-based methods found significant associations for the less specific gene set *T-Cell Differentiation*, but not for *Th1 & Th2 differentiation.* Thus, subtle immune effects may be more readily apparent from the row variances of **M**_1_ than from the methylation associations obtained by adjusting for the reference-based cell proportions.

### Additional Analyses on Datasets with Normal and Pathological Tissue

Finally, in an analysis of cell proportions **Ω** obtained using the Roadmap-derived methylomes as a reference, notable distinctions between normal and pathological tissues were revealed (Figures S9.1 through S9.3). In particular, gastric tumors differed from normal gastric tissues in having greater immunological/inflammation content but lesser gastrointestinal content, atherosclerotic carotid (and to some extent atherosclerotic aorta) differed from normal aorta in having greater immunological/inflammation content but lesser muscular content, and cirrhotic tissues differed from normal liver tissues in having greater immunological/inflammation content but lesser gastrointestinal content (with the pattern more striking for cirrhotic tissues related to viral infection than for cirrhotic tissues related to alcohol abuse).

## Discussion

We have proposed a simple method for reference-free deconvolution that provides both interpretable outputs, i.e. proportions of putative cell types defined by their underlying methylomes, as well as a method for evaluating the extent to which the underlying reflect specific types of cells. We have demonstrated these methods in a wide array of methylation datasets in various tissues and focused on differing exposures or outcomes.

Overall our deconvolution approach is similar to many others that have been proposed^29-33^. In particular, it is very similar to a recent publication that applied a convex-mixtures approach to deconvolve RNA expression^33^. Our approach differs from this one in that it deconvolves DNA methylation, with a corresponding constraint on the values of **M**; in addition, importantly, we have provided a more comprehensive approach for interpreting the resulting columns of **M**.

We have provided a novel approach for estimating the number 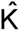 of cell types, which we have shown to reflect the level of cellular heterogeneity anticipated from each tissue we analyzed.

More heterogeneous tissues (blood, breast, and gastric tissues) resulted in higher values of 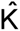, while the more homogeneous tissues had lower values: 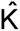 = 1 for the (admittedly small) isolated lymphatic and blood vessel endothelium data sets, while 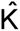 = 2 for sperm and umbilical cord endothelial tissues. Note, however, that Figure 2b reflects ambiguity in the choice of 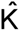 for sperm and equally supports the choice 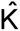 = 1; similar plots shown in Figure 2b unambiguously suggest 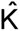 > 1 for two blood data sets and for an artery data set. A similar plot for UV-as (not shown) displayed an unambiguous preference for the choice of 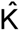 = 2, but the two putative types of cells did not associate with any metadata (Table 2). Taken together, these results demonstrate that our algorithm returns reasonably reliable values of 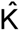 reflecting cellular heterogeneity.

Cell mixture proportions **Ω** were typically significantly associated with major phenotypes of interest, with the notable exception of sperm, umbilical vein endothelium, and placental artery (the former two assumed to be homogenous tissues). Thus, the radically dimension-reduced DNA methylation data in **Ω** can still retain strongly significant associations with major phenotypes of interest. However, for other covariates, especially demographic confounders, there was considerable variation in significance, demonstrating that **Ω** can also show null associations with some covariates. Taken together, the results show that **Ω** can distinguish signal from noise. As the limma analysis demonstrated, residual signal can still exist in **Y** even after adjusting for **Ω**, although often in a more diminished capacity. In a few rare cases, the signal increased after adjusting for **Ω**. Taken together, these results suggest that a substantial proportion of the association between **Y** and phenotypic metadata **X** can be factored through the decomposition **Y** = **MΩ**^T^, occasionally clarifying the residual signal, but more often diminishing it. This finding is significant, as it would strongly suggest that the results of the vast majority of EWAS studies are driven by physiologic changes of the underlying composition of cells within the samples obtained. This is nicely highlighted by a recent report identifying a specific cell type driving the associations between smoking and changes in DNA methylation in peripheral blood^23^. This is in contrast to the prevailing current interpretation of most findings, which has aligned more strongly with the concept of metastable epialleles. These alleles represent loci where environmental conditions during development dictate ‘setpoints’ for the levels of methylation at particular gene sequences that are consistent across tissues within any person, also yielding differences in concordant gene expression^45-48^. The methods described in our work may have some utility for future discovery of these alleles in that within-person, cross-tissue comparisons of methylation profiles would be expected to be enriched for metastable alleles when the loci that are reflective of subsets of cell types are described and removed from comparisons.

On the other hand, we demonstrated that columns of **M** correlate with external biological annotation data in a manner concordant with their interpretation as methylomes specific to constituent cell types. *Blood DMP* status showed the highest associations in blood data sets, although also somewhat high associations in *L[np]* and *AR[np]*, data sets having tissues with potentially inflammatory components to pathology. Data sets with tumors *(BR-tcga[t], br-1[t], br-2[t], br-3[t], g[nt]*, and *g[t])* demonstrated high associations with *PcG Loci*, reflecting the mitotic activity of tumors; that normal gastric tissue, *g[n]* also showed a high association with *PcG Loci* is consistent with the high level of cellular turnover in gastric tissues. The Bilenky DMRs showed the strongest associations for breast tissues, consistent with the fact that the Bilenky DMRs were obtained from breast tissue, but also demonstrated strong associations for liver pathologies, and in arterial tissues *AR[n]* and *AR[np].* Breast and arterial tissues have a mix of epithelial and smooth muscle tissues, which may explain the arterial results. The association in *L[np]* may reflect the fibrous character of pathological liver tissue. REMC DMRs demonstrated only weak correlation or strong negative correlation with all tissues, perhaps reflecting the embryonic/developmental nature of the REMC DMRs.

Comparisons with DMPs constructed from Roadmap WGBS data also demonstrated that the columns of **M** reflect epigenetic content concordant with anatomical expectations; in particular, blood datasets displayed associations with DMPs distinguishing CD34+ hematopoietic stem cells vs. thymus or spleen, as were datasets *L[np]* and *AR[np]*, which both included tissue pathologies involving inflammation and immune response. Arterial data sets displayed associations with DMPs distinguishing smooth muscle from endothelium. Associations with Roadmap-based DMPs were typically weak for homogeneous tissues, in particular sperm. Interestingly, the normal liver tissue data set *L[n]* had mostly weak negative associations; one possible explanation is that the primary tissues available from the Roadmap were too dissimilar from normal liver tissue to distinguish subtle anatomical features. Using the Roadmap data as a pseudo-reference, normal and pathological tissues were revealed to differ anatomically along anticipated lines, specifically in that pathological tissues had greater cellular content reflective of immune or inflammation processes, and lesser gastrointestinal content (gastric and liver tissue) or muscular content (arterial tissue). Taken together, these results suggest that the columns of **M** reflect methylomes of constituent cell types.

We remark that the unsupervised deconvolution approach we have proposed cannot be guaranteed to recover the methylomes of all constituent cell types; instead, it recovers the major axes of cellular variation. This was evident in the comparison of reference-based and reference-free deconvolution of blood datasets *BL-ra* and *BL-as*; for these datasets, the reference-free approach recovered the linear combination of reference methylomes most relevant to characterizing the underlying variation. However, when “re-mixed” back to proportions of known cell types using a reference methylome, associations with phenotypic metadata were consistent with those obtained from reference-based deconvolution. While on its surface this suggests that the reference-free approach has no value when a reference methylome is known (as is the case with blood), further analysis of the residual information in the unsupervised deconvolution demonstrated that reference-free deconvolution may reveal distinctions in cell type relevant to characterizing the underlying variation in the dataset but more subtle than the potential distinctions fixed in advance by the reference set. For example, the unsupervised approach identified two strongly significant processes in the rheumatoid arthritis data set, *Th1 & Th2 differentiation* and *T-Cell Polarization*; this finding is consistent with known Th1/Th2 differentiation processes^49, 50^ and T-Cell polarization process^51, 52^ involved in the etiology of rheumatoid arthritis. Similarly, the unsupervised approach identified *Regulators of T-Cell Activation* as a significant process in the dataset investigating arsenic exposure in Bangladesh. In fact, the impact of arsenic exposure on regulation of T-cells has been noted^53^, and even observed in another study conducted in Bangladesh^54^. We remark that we obtained somewhat different results using reference-based cell proportions **Ω**_0_ in *limma* to adjust for cell type. Thus, the reference-free approach may provide important information that complements a reference-based approach.

As a general point, we have demonstrated the links suggested in Figure 1; thus, we have shown that it is possible to use a reference-free approach to characterize the extent to which phenotypic associations with DNA methylation data are explained by differences in constituent cell types. We remark that such distinctions may be subtle, such as variation in smooth muscle content or the presence of leukocytes with specialized immunological states. There may still exist associations residual to those with variations in putative underlying cell types, although they will often be diminished after adjusting for cell type in the manner we have proposed. Other reference-free approaches can also distinguish between associations driven by variation in cell type and those that are more focal to individual CpG sites, but our proposed method has several advantages over these existing methods. The first is that it is not particularly intensive computationally; the second is that it provides an easy and interpretable way to estimate the underlying number of constituent cell types; the third is that it provides estimates of cell-proportions that are directly interpretable and comparable with estimates obtained from reference data sets. In particular, our method provides a means for extracting information that is more subtle than that available from reference data sets but may nevertheless reflect additional variation in constituent cell types. While similar insights may be obtained simply by examining the CpG-specific associations, we note that there is ongoing controversy on what “adjustment for cell-type” means in the context of EWAS analysis. We have previously argued that all epigenetic variation is ultimately mediated by cell-type, if the meaning of “cell-type” is conceived of broadly enough^20^; a more useful framing of the question is how to identify types of cells that are relevant to the biological variation being studied. Our proposed approach helps in partitioning the underlying variation into units that resemble cell-specific methylomes, so that these methylomes or the overt functional characteristics of these cells may be further analyzed using additional biological characterization data.

We remark on a few current limitations of our approach. One is that we have used a crude gene-set procedure based on variance, which removes “signed” information and thus precludes the use of algorithms based on expression signature, such as CTen^55^. Another related limitation is a lack of relevant annotation data. Further work is necessary to adapt the method we have proposed here to “signed” comparisons, thus enabling a wider array of annotation tools, and to develop other relevant annotation datasets relevant to identifying subtle cell types.

## Methods

### Empirical Examination of Proposed Methods

We removed chromosome Y data from all datasets; and we also removed chromosome X data from all but the breast datasets. For the 450K data sets downloaded from TCGA, and for the 450K data collected to investigate associations with arsenic exposure in Bangladeshi neonates^3, 9^, we used the *FunNorm* algorithm *(Bioconductor* package *minfi)* to process the raw *idat* files; we obtained all other data sets as processed average beta values from Gene Expression Omnibus (GEO). For 450K data sets, we excluded CpGs with cross-hybridizing probes or probes with SNPs^56^, and used the *BMIQ* algorithm^57^ *(Bioconductor* package *wateRmelon)* to align the scales of Type I and Type II probes. Finally, for each data set, we excluded CpGs having missing measurements for over half the specimens.

### Associations with Phenotypic Metadata

As described in Supplementary Section S4, permutation tests were used to assess omnibus significance of covariates **X** with fitted cell proportions **Ω**. As described in Supplementary Section S5, we further compared associations of **Y** with **X** before and after including terms from **Ω** in the regression model for **Y**, using the *limma* procedure^58^ (via the R package *limma*) to compute regression coefficients, using the R package *qvalue* to estimate both q-values and the overall proportion **π**_0_ of null associations.

### Interpretation of Putative Cell Types

We obtained a list of DMPs for differentiating distinct major types of leukocytes *(Blood DMPs)* from the Reinius reference set^25^, and constructed a set of CpGs mapped to genes considered Polycomb Group proteins *(PcG loci)*, compiled from four references^59-62^ as in our previous articles^20, 27^. We also constructed a set of CpGs based on differentially methylated regions (DMRs) obtained from WGBS data collected by the Epigenomics Roadmap Project. Supplementary Section S6 describes the details of the construction of these DMP sets. In addition, we developed a novel approach based on WGBS data from the Roadmap Epigenomics Project for 24 primary tissues, described in detail in Supplementary Section S7.

### Additional Analyses on 450K Blood Datasets

To compare reference-based analysis with our proposed approach, we analyzed the two 450K blood data sets, *BL-ra* and *BL-as*, estimating for each data set two sets of cell-type proportion matrices (*K* = 7): **Ω**_0_ (reference-based) and **Ω**_1_ (reference-free). Details appear in Supplementary Section S8. Briefly, to obtain the mixing matrix **Ψ** that relates matrices **M**_0_ and **M**_1_, we used a constrained projection similar to that used to obtain the reference-based cell proportion matrix **Ω**_0_, and compared **M**_1_ with **M**_1_ – **M**_0_**Ψ**^T^ by identifying CpGs with high variation across their constituent methylomes. In addition, we compared these highly varying CpGs with with immune activation and immune regulation pathways compiled from six sources^63-69^.

### Additional Analyses on Datasets with Normal and Pathological Tissue

We projected Infinium data from each of the three datasets sets *g[nt], AR[np]*, and *L[np]* onto the profile matrix **M** obtained by decomposing the Roadmap WGBS data (**Y** = **MΩ**^T^); we then averaged the resulting specimen-specific cell proportions **Ω** over tissue status (normal gastric tissue vs. gastric tumor, normal aorta vs. atherosclerotic aorta and atherosclerotic carotid, and normal liver vs. alcohol-related cirrhotic liver and cirrhotic liver due to viral infection). Details and results appear in Section S9.

## Acknowledgements

Funding for this work was provided by grants from the National Institutes for Health: NIMH R01MH094609 (EAH and CJM), NIEHS P01 ES022832 (CJM), K01 ES017800 (MLK), R01ES024991 (EAH, TAI), R01ES015533 (DC). Funding was also provided by EPA grant RD83544201 (CJM).

